# The Secretomes of Painful Versus Nonpainful Human Schwannomatosis Tumor Cells Differentially Influence Sensory Neuron Gene Expression and Sensitivity

**DOI:** 10.1101/734699

**Authors:** Kimberly Laskie Ostrow, Katelyn J. Donaldson, Michael J. Caterina, Allan Belzberg, Ahmet Hoke

## Abstract

Schwannomatosis is a multiple tumor syndrome in which patients develop benign tumors along peripheral nerves throughout the body. The first symptom with which schwannomatosis patients often present, prior to discovery of tumors, is pain. This pain can be debilitating and is often inadequately alleviated by pharmacological approaches. Schwannomatosis-associated pain can be localized to the area of a tumor, or widespread. Moreover, not all tumors are painful, and the occurrence of pain is often unrelated to tumor size or location. We speculate that some individual tumors, but not others, secrete factors that act on nearby nerves to augment nociception by producing neuronal sensitization or spontaneous neuronal firing. We created cell lines from human SWN tumors with varying degrees of pain. We have found that conditioned medium (CM) collected from painful SWN tumors, but not that from nonpainful SWN tumors, sensitized DRG neurons, causing increased sensitivity to depolarization by KCl, increased response to noxious TRPV1 and TRPA1 agonists and also upregulated the expression of pain-associated genes in DRG cultures. Multiple cytokines were also detected at higher levels in CM from painful tumors. Taken together our data demonstrate a differential ability of painful versus non-painful human schwannomatosis tumor cells to secrete factors that augment sensory neuron responsiveness, and thus identify a potential determinant of pain heterogeneity in schwannomatosis.

## Introduction

Patients with schwannomatosis (SWN) develop multiple tumors along major nerves of the body^1^. In most cases the tumor burden is so great that they cannot be removed by surgical intervention ^2^. Sixty-eight percent of SWN patients also report chronic pain, which is often debilitating ^3^. Pain in SWN has been described as localized in some patients and diffuse in others ^4^. The size and location of the tumor(s) do not necessarily relate to the severity of pain experienced by the patient ^5^. Thus, while surgical removal of an individual tumor may provide pain relief in some cases, the surgeon is challenged with which tumor to remove. Additional confounding factors (e.g., nerve deficits, tumor recurrence, return of pain) may also affect surgical outcome ^6,7^. Quality of life in Schwannomatosis patients is predominantly affected by pain, especially in those with a large tumor burden. Therefore, treatment approaches other than tumor-focused surgery needs to be developed.

Neuropathic, nociceptive, and inflammatory types of pain have all been reported in SWN. The pain described by SWN patients can vary from burning to pain that feels like electric shocks ^8,9^. Yet, the molecular mechanisms by which some SWN tumors, but not others, elicit unremitting pain are unknown ^3^. Given the close interactions between Schwann cells and neurons, we have chosen to study the potential role of Schwann cells in neuropathic pain in schwannomatosis. It is well documented that in models of peripheral nerve injury, Schwann cells become activated and release soluble pro-inflammatory cytokines as well as other chemoattractants^10–12^. These soluble substances recruit macrophages, induce myelin clearance, and promote nerve regeneration. Many pro-inflammatory cytokines (e.g., TNF-alpha, IL-1α, IL-1β, Il-6, CCL2/MCP-1, CCL3/MIP-1α and GM-CSF) are upregulated post-injury and many of these can sensitize nociceptors ^10,13–15^. Anti-inflammatory cytokines such as IL-10 are also produced to counterbalance the pro-inflammatory cytokines ^16^, since a regulated inflammatory response is necessary for nerve repair and regeneration.

Here, we examine the hypothesis that painful SWN cells secrete substances, such as cytokines, into the extracellular space that sensitize neurons and make them easier to excite. Knowledge gained from this study will help pinpoint candidate molecules and pathways that can be targeted for drug development for the treatment of schwannomatosis-related pain.

## Results

To test our hypothesis outlined above, and to define the mediators that drive pain in some patients with SWN, we have been examining the “secretome” of SWN tumor cells in culture. Schwannomatosis cell lines were established from fresh patient tumors from surgical excision. On the day of surgery we asked patients to rate their pain on a scale from 1 to 10 and categorized them as such: Pain score (PS) 0 no pain, 1-3 low pain; 4-6 moderate pain; 7-10 severe pain. We have established 7 cell lines from SWN patients with varying degrees of pain. Three were non-painful or had a pain score < 3, one has moderate pain (PS=6) and three reported high pain scores (PS 7-9). All cells were immortalized with SV40 and demonstrated S100 positivity ^17^. Schwannoma conditioned media was collected after 48 hours and frozen for subsequent experiments.

### Effect of CM on neuronal sensitivity in vitro

To determine the effects of the schwannoma secretome on sensory neurons, primary wild-type mouse dorsal root ganglion (DRG) neurons were incubated in CM derived from (1) painful SWN cells, (2) non-painful SWN cells, and (3) normal human Schwann cells. After a 48-hour incubation, the DRG neurons were tested for their ability to respond to an ascending series of KCl concentrations, using fura-2 based Ca2+ imaging. No visible differences in neurite sprouting could be observed in DRGs treated with painful versus non-painful CM (Supplemental figure 1). Neuronal depolarization in response to KCl or other excitatory stimuli triggers action potentials, which in turn triggers Ca2+ influx through activation of voltage-gated Ca2+ channels. Neuronal stimulation can also trigger Ca2+ release from intracellular stores. Ratiometric Ca2+ imaging is therefore used as an indirect measure of neuronal activation at a given strength of stimulation ^18–21^. We observed that neurons treated with media conditioned by painful schwannomas were more responsive to KCl than those treated with medium conditioned by the non-painful schwannomas or normal Schwann cells (Figure 1). In neurons treated with CM from non-painful schwannoma cells or normal human Schwann cells, Ca2+ responses were reliably observed at 15 mM KCl and increased with KCl dose. By comparison, KCl evoked responses were larger in cells treated with CM from painful schwannoma cells, with more neurons responding at the lowest KCl dose (10mM) and a greater area under the curve at each KCl dose. The effect of painful CM was reduced when it was heated to 80° C for 30 minutes prior to DRG treatment, suggesting that a thermolabile component, most likely protein(s) secreted by the tumor, are contributing to neuronal sensitization (Supplemental Figure 2).

**Figure 1:**
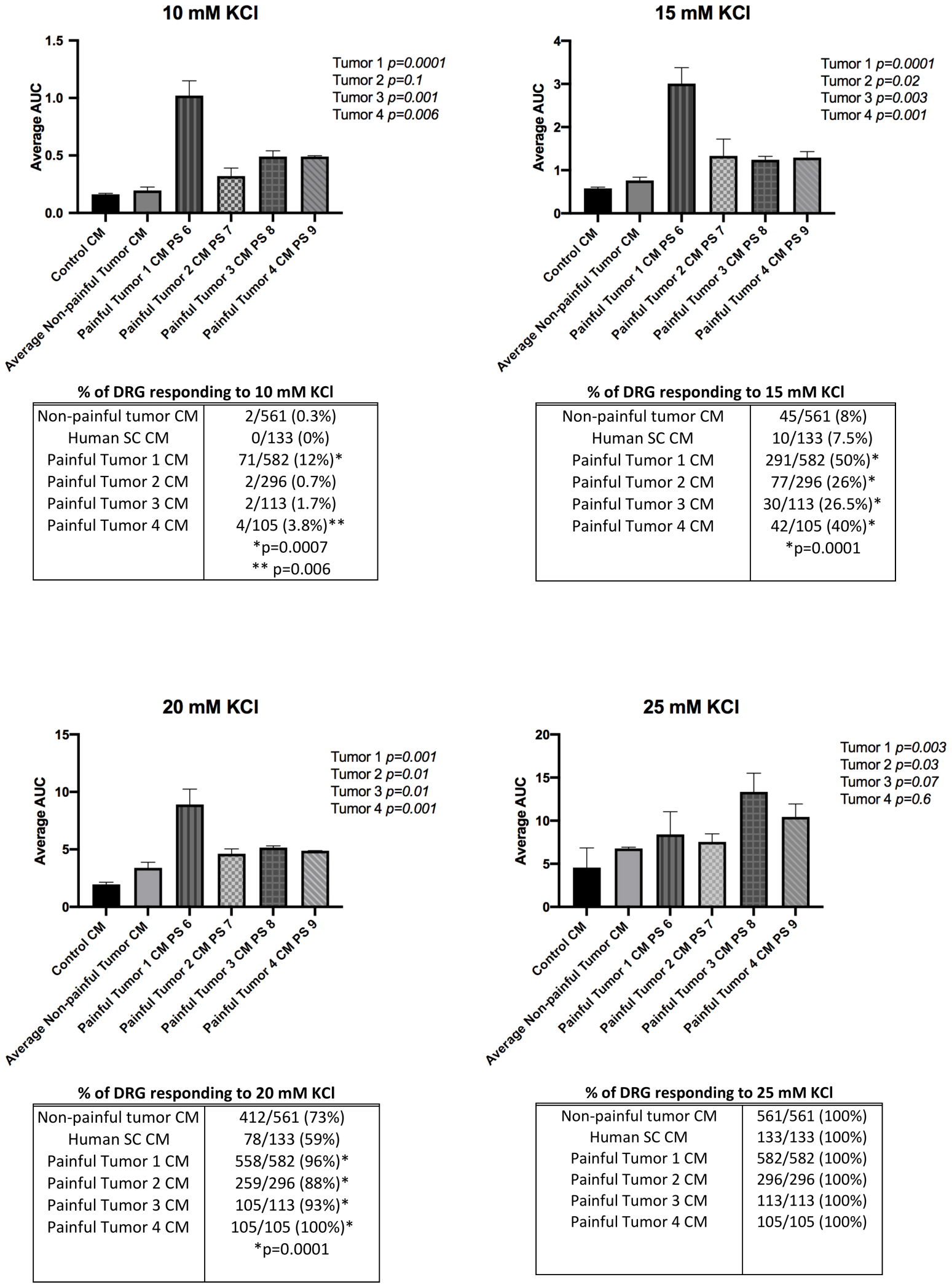
Primary mouse DRG neuron cells dose-response to KCl after pre-treatment with schwannoma conditioned media. DRG cells were pre-treated with painful schwannoma CM, non-painful schwannoma CM, or normal Human Schwann cell CM for 48 hours prior to imaging. 10 coverslips containing ~30/slip were tested for each condition. DRG cells were perfused with increasing amounts of KCl (10mM KCl, 15mM KCl, 20 mM KCl, 25mM KCl). Each perfusion was 30 seconds in duration with a washout with 3mM KCl for 2 minutes prior to the next increasing dose. Fura-2 ratio measurements were recorded at 2 second intervals. The datapoints on the graph were corrected for differing baseline readings of Fura-2. The graph represents the average fura-2 ratio across all coverslips for each of the CM treatments. Maximum response occurred 30 seconds after each dose was administered. Fura-2 ratio returned to baseline after 60 seconds from the start of the dose. The dose response was similar for non-painful schwannoma CM pre-treatment and Human Schwann cell CM pretreatment. Area under the curve was calculated for each KCl treatment. The effect of KCl started immediately after treatment, peaked at 30 seconds, and lasted for 60 seconds (Supplemental Figure 2). A significantly larger effect (p<0.03) at each dose (10 mM-20mM KCl) was demonstrated in the cells pre-treated with painful schwannoma CM as demonstrated by larger area under the curve. For ease of presentation, data for the non-painful tumors (n=3) were averaged together. The differences in AUC between non-painful CMs were not significant (Supplemental figure 2). ANOVA analysis with Dunnet multiple comparisons was used to compare the effects of CM groups (Graphpad Prism8). The percentage of cells responding to KCl increased in a dose dependent manner. CM from painful SWN caused a larger percentage of cells to respond to KCl. An individual cell’s response to stimuli was considered to be positive if the fura-2 ratio was greater than 0.1 after baseline subtraction. Chi-squared analysis was performed to test significance between the percentage of cells responding to painful and non-painful CM.

In addition, whereas all neurons responded to 25 mM KCl regardless of the CM with which they were pretreated, a larger percentage of neurons responded to 15 mM and 20 mM KCl when pre-treated with CM from painful SWN tumors (Figure 1). Even at the lowest dose of KCl (10 mM), neurons pre-treated with CM from painful tumor 1 showed not only a significantly larger response (AUC 1.0 vs. 0.19 p=0.0001) but also a larger percentage of cells responding (12% of DRG neurons responded, p<0.0001, Chi-squared) when compared with the non-painful groups. Together, these findings indicate that exposure to CM from painful SWN tumor cells results in greater sensory neuron responsiveness to depolarization than does exposure to CM from non-painful SWN tumor cells or control Schwann cells.

### Effect of painful schwannoma conditioned media on DRG neuronal expression of pain-related genes

Primary mouse DRG neurons were treated with schwannoma CM for 48 hours as described above. DRG neurons were then harvested and RNA isolated. The Qiagen RT2 profiler array for neuropathic and inflammatory pain was used to assess gene expression changes by qPCR. Several genes related to inflammatory pain were upregulated in DRG neurons after treatment with CM from a painful schwannoma, relative to nonpainful CM (Figure 2). Common to all painful tumor CM treatments was the upregulation of Bradykinin receptor by the DRG neurons. Some heterogeneity in gene expression was observed in DRGs treated with painful tumor CM. The nociceptive ion channel transient receptor potential ankyrin (TRPA1) was up-regulated in DRGs by several painful CM samples (Painful Tumor 2 p=0.04, Painful Tumor 3 p=0.05) as was interleukin 1β (Painful Tumor 1). CCKBR was upregulated in DRGs treated with CM from painful tumor 4.

**Figure 2:**
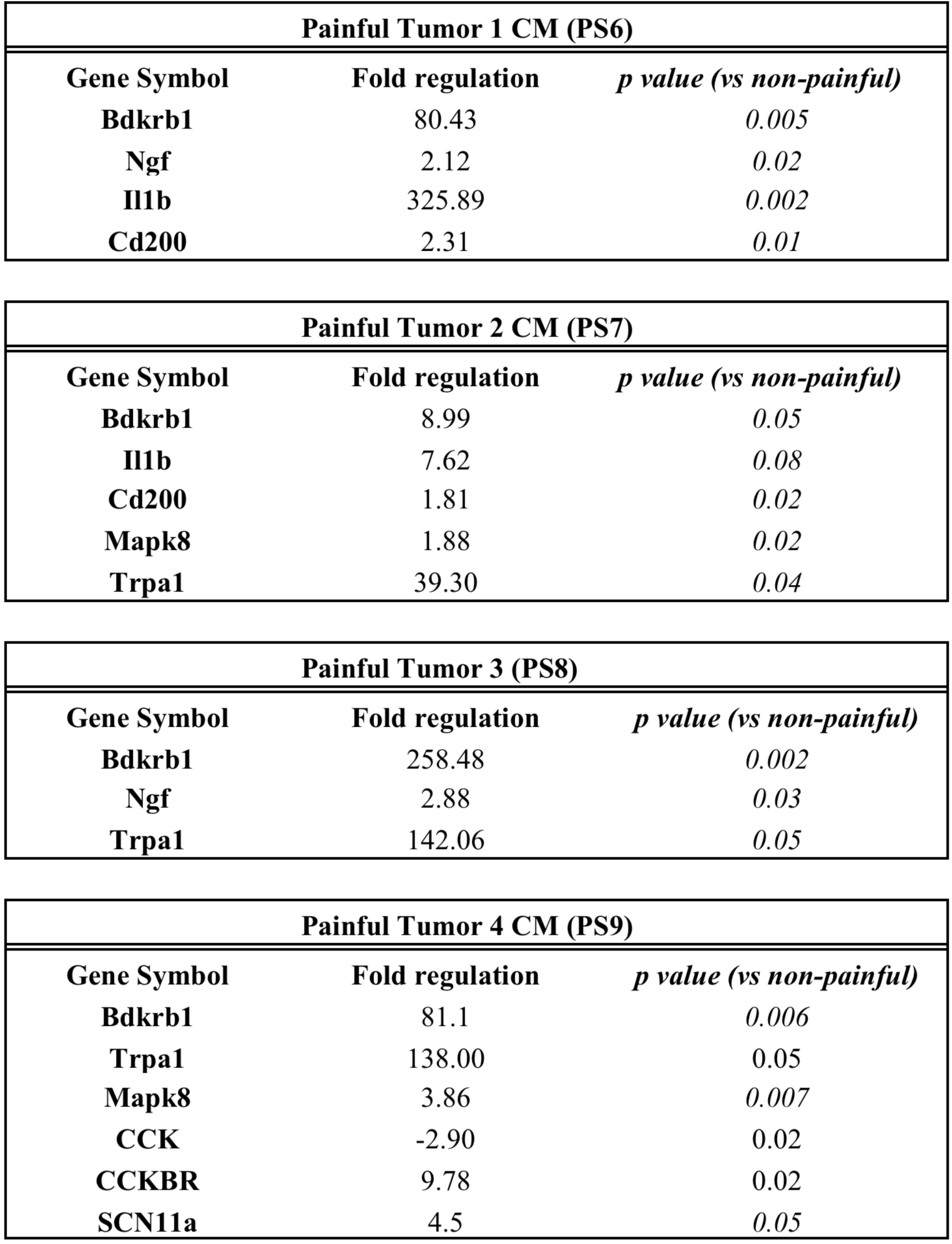
Painful SWN tumor CM influences transcription of pain-related genes in DRG neurons. Primary mouse DRG neurons were treated with painful or nonpainful schwannoma CM for 48 hours as described above. RNA was isolated from the neurons and subjected to qRT-PCR using the Qiagen RT2 profiler array for mRNAs related to neuropathic and inflammatory pain. Of the 84 pain-related genes represented in this array, 64 could be detected in untreated mouse DRG neurons. Among these, a number of genes appeared to be differentially expressed in neurons after treatment with painful vs. nonpainful schwannoma CM. Genes demonstrating a 1.5 fold difference in regulation and a p value <0.05 was considered significant. For qRT-PCR, Fold change in gene expression was determined by the 2^(- Delta Delta Ct) method. Statistical significance of the RT2 PCR data was determined using a Student’s t-test (https://dataanalysis.qiagen.com/pcr/arrayanalysis). Differences in gene expression were also noted between cells treated with different painful tumor CMs.

### Heterogeneous cytokine secretion by painful and non-painful SWN tumor cells

We performed protein-level analysis of cytokines secreted into CM collected from painful vs. non-painful SWN cell lines using the Human XL Cytokine Array (*R&D Systems*). We observed multiple differences in levels of cytokines secreted by painful and non-painful SWN tumors (Figure 3). Specifically, GDF-15/MIC-1 Endoglin, GM-CSF, CD147. CXCL1, PTX3 and were found at higher levels in CM from painful tumors. We also observed heterogeneity in cytokine secretion between painful tumors. Several painful tumors secreted higher levels of IL8, VEGF, CCL5/RANTES, and IL6, compared to the non-painful tumors (Figure 3).

**Figure 3:**
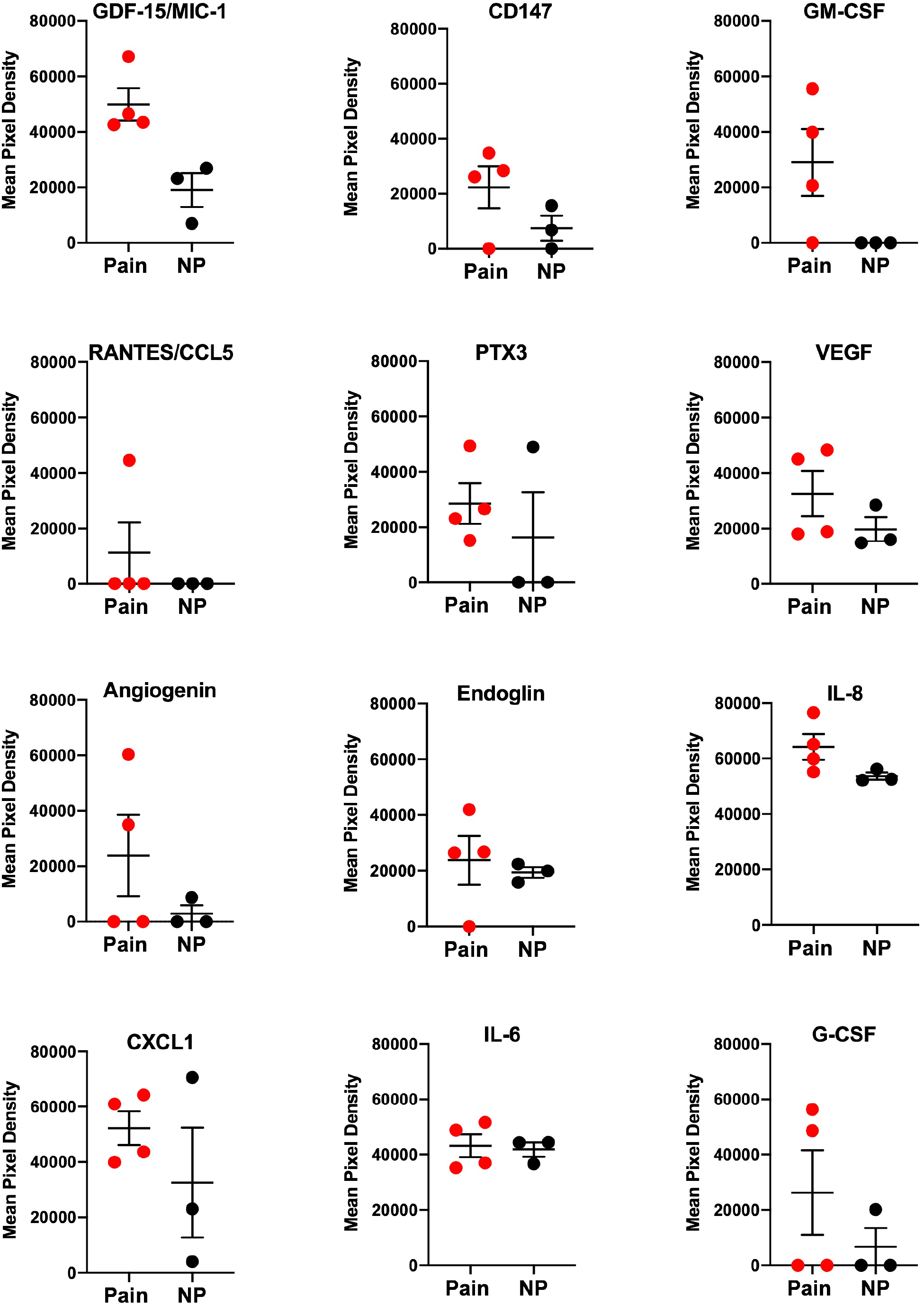
Elevated levels of secreted cytokines are present in conditioned media from painful schwannomatosis cell cultures. The Human Cytokine XL array (R&D Systems) detected the presence of cytokines in conditioned media from schwannomatosis cultures. Proteome blots were exposed to X-ray film for 2 minutes. Quantification of cytokine level was determined by pixel density of the proteome blots using ImageJ. Internal controls (3 positive and 1 negative) are used to normalize differences in total protein input (Supplemental Figure 7). Three non-painful tumors (black circles) were compared to 4 painful tumors (Red circles). Mean pixel densities are shown for each sample. Horizontal line is mean +/− SEM

### Effect of TRPA1 and TRPV1 activators on DRGs after pre-treatment with tumor CM

Upon analysis of the secreted cytokine/chemokine profiles in tumor CM and changes in gene expression in DRG neurons after treatment with tumor CM we are narrowing in on candidate pain pathways. We observed TRPA1 overexpression in DRG neurons that were treated with CM from painful tumors also secreted potential activators of TRPA1 (CCL5/RANTES and GM-CSF)^22,23^

We therefore hypothesized that TRPA1 is involved in pain signaling in painful SWN tumors. We treated mouse DRG neurons with painful tumor CM or non-painful tumor CM for 48 hours, then performed Ca2+ imaging during challenge with the TRPA1 agonist cinnamaldehyde (200 μM). Overall, DRG cells treated with SWN CM showed an increased sensitivity to cinnamaldehyde but the magnitude of response to cinnamaldehyde was significantly greater in cells pre-treated with painful tumor 3 CM (painful tumor 3 PS 8 AUC=29 vs. nonpainful tumor AUC=14; p=0.03; Figure 4). The percentage of cells responding to cinnamaldehyde also increased in DRG cells pretreated with CM from painful tumor 2 and 3 with statistical significance (non-painful tumors 17% responded to cinn vs. painful CM 2 39% responded, painful CM 3 28% of DRGs responded p=0.001) (Figure 4). Both of these painful CM samples also caused increased transcription of TRPA1 in DRG neurons (Figure 2).

**Figure 4:**
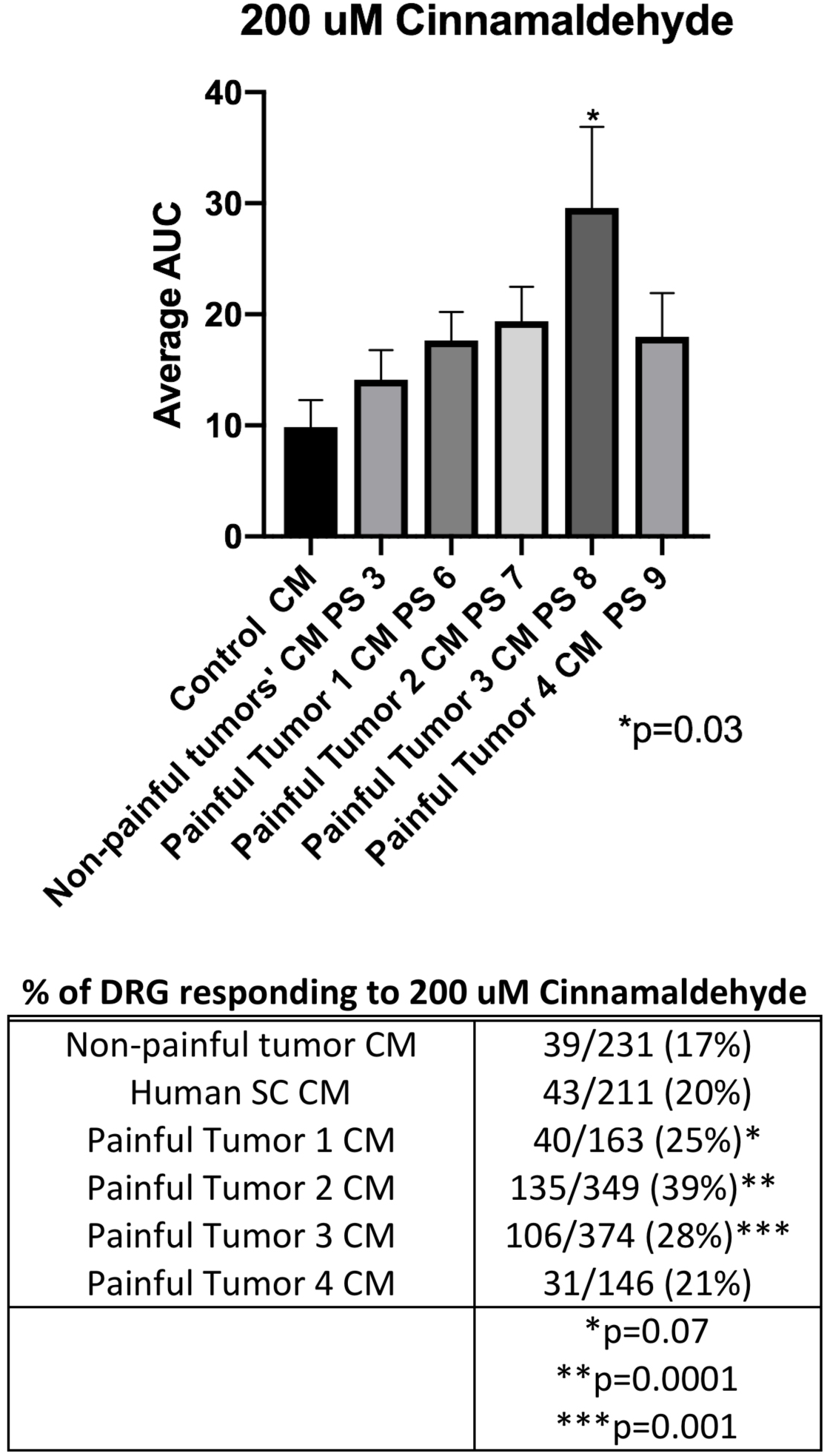
Conditioned media from painful SWN tumors sensitize DRG neurons to the TRPA1 agonist cinnamaldehyde. DRG cells were pre-treated with painful schwannoma CM, non-painful schwannoma CM, or normal Human Schwann cell CM for 48 hours prior to imaging. DRG cells were perfused 200 uM Cinnamaldehyde for 120 sec with a washout with 3mM KCl buffer for and additional 2 minutes. Fura-2 ratio measurements were recorded at 2 second intervals. The datapoints were corrected for differing baseline readings of Fura-2. Area under the curve was calculated for each treatment. The effect of cinnamaldehyde started 30-60 seconds after treatment peaked at 120 seconds, and lasted for 4 minutes. Painful tumor 3 CM increased the responsiveness of the DRGs to cinnamaldehyde as calculated by area under the curve p=0.03. One way ANOVA with multiple comparisons was employed to determine statistical significance. The percentage of DRG cells responding to cinnamaldehyde increased after pre-incubation with CM from painful SWN tumors 2 and 3. Chi-square analysis was used to compare each painful tumor CM treatment to non-painful tumor CM treatment. p<0.05 is considered significant.

While not up-regulated at the transcriptional level we examined the potential role of the nociceptor TRPV1 in SWN-related pain. TRPV1 is a sensor for noxious heat, pH, and its most recognized agonist, capsaicin ^24,25^. It is expressed in DRG neurons and is known for its role in hyperalgesia. One of our candidate SWN secreted cytokines, GM-CSF sensitizes neurons to capsaicin^26^. Therefore, using the same methodology as used for cinnamaldehyde described above, we pretreated DRG cells with painful or nonpainful CM for 48 hours and tested responsiveness to sub-saturating capsaicin. All DRG cells pre-treated with CM from painful tumor cells had a larger response to 25 nM capsaicin than did cells pre-treated with CM from non-painful tumor cells or control cells, as demonstrated by either an increased magnitude of response (painful tumor 3 CM AUC=15.9, painful tumor 4 CM AUC=16 vs. non-painful tumor AUC=7.0; p=0.0001) or an increase in the percentage of cells responding to capsaicin (44% CM1, 41% CM2, CM4 65% vs 29% non-painful) (Figure 5).

**Figure 5.**
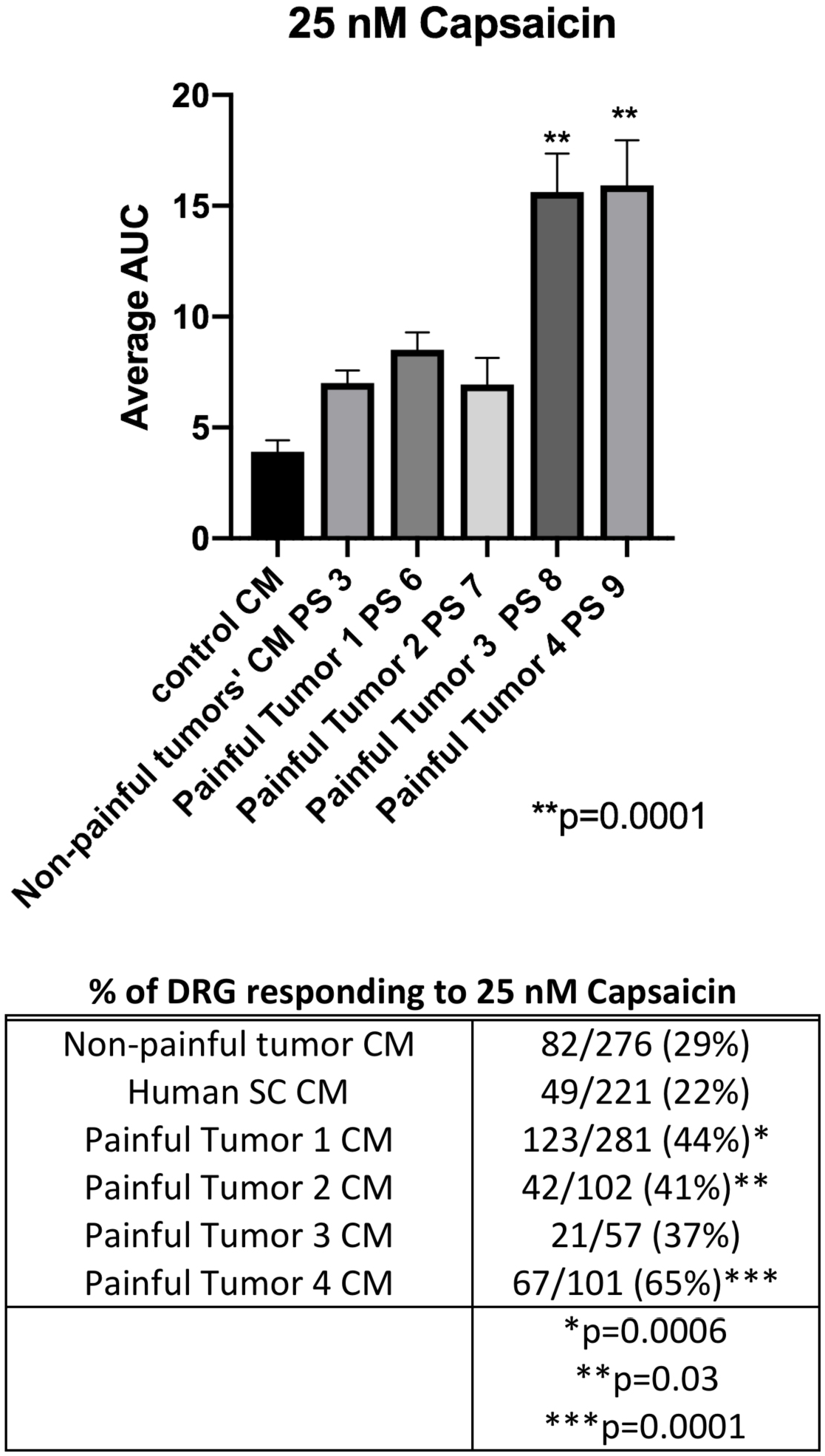
Conditioned media from painful SWN tumors sensitize DRG neurons to the TRPV1 agonist capsaicin. DRG cells were pre-treated with painful schwannoma CM, non-painful schwannoma CM, or normal Human Schwann cell CM for 48 hours prior to imaging. DRG cells were perfused 25nM capsaicin for 30sec with a washout with 3mM KCl buffer for 2 minutes. Fura-2 ratio measurements were recorded at 2 second intervals. The datapoints were corrected for differing baseline readings of Fura-2. Area under the curve was calculated for each treatment. The effect of capsaicin started immediately peaked after 30 seconds lasted for 1 minute (Supplemental Figure 5). CM from painful tumors 3 and 4 CM increased the responsiveness of the DRGs to capsaicin as calculated by area under the curve p=0.0001. One way ANOVA with multiple comparisons was employed to determine statistical significance. The percentage of DRG cells responding to capsaicin increased after pre-incubation with CM from each painful SWN tumors. Chi-square analysis was used to compare each painful tumor CM treatment to non-painful tumor CM treatment. p<0.05 is considered significant.

While the proteins secreted and the effect on DRG neurons are heterogeneous between painful tumors, these data support our hypothesis that painful tumors secrete proteins that sensitize neurons *in vitro*.

## Discussion

The mechanisms driving the pain phenotype in SWN patients are unknown. Patients vary in their presentation of pain and describe their experiences with pain as acute, chronic, localized, or widespread ^3^. Pain in SWN patients is classified as inflammatory or neuropathic. The drugs that are available to treat pain are grossly ineffective in this patient population. Currently the standard of care to alleviate pain is to remove the tumor that is thought to be the cause of pain ^5^. This is a difficult undertaking. The surgeon must attempt to localize the tumor that is causing pain. When a patient has a moderate tumor burden and pain is widespread there is uncertainty as to which tumor(s) should be removed. Even if the tumor is removed, the pain may not be resolved or may recur if the tumor grows back ^5–7^. This results in a viscous cycle of multiple surgeries to hunt down the source of pain. Therefore, alternative strategies for pain therapy must be investigated.

The reasons why some SWN tumors cause significant pain and others don’t remain a mystery. Given this heterogeneity, we hypothesized that painful tumors might secrete specific factors that act on the nearby sensory nerves, causing the latter to be sensitized. To test our hypothesis, we isolated CM from SWN cell lines that were developed in our lab. Cell lines were created from tumors surgically excised from SWN patients who had tumors removed due to pain and others that did not cause pain but were excised for another reason ^17^. CM from 4 painful SWN tumors and 3 non-painful tumors were collected for analysis. Each CM sample was tested for the ability to increase the sensitivity of DRG neurons to painful stimuli, assessed for the ability to alter gene expression in DRG neurons, and probed for elevated levels of pro-inflammatory cytokines. By examining the data we are beginning to see a clearer picture of the key pathways involved in schwannomatosis-related pain.

Ratiometric Ca2+ imaging was used as an indirect measure of neuronal activation ^18–21^. Small diameter DRG neurons (10-20 micron diameter) were sensitized to conditioned media isolated from painful schwannoma tumors. We observed a shift to lower KCl concentrations required for responses as well as larger integrated response sizes (AUC) at all KCl concentrations after pre-treatment with CM from painful tumors as opposed to non-painful tumors CM or CM from normal human Schwann cells (Figure 1). These data indicate that CM from painful tumors renders sensory neurons more responsive to depolarization than does CM from non-painful tumors or normal human Schwann cells but do not identify the roles of pain specific pathways.

We asked whether the effect of painful tumor-derived CM on DRG neuron sensitivity was associated with changes in transcriptional regulation of neuronal genes relevant to pain. Significant differences in gene expression were detected in DRG neuronal cultures after treatment with CM isolated from painful tumors, compared with CM from non-painful tumors. Several genes related to inflammatory pain were highly upregulated in the DRGs treated with CM from painful tumors, and their encoded proteins thus constitute downstream candidates for further investigation in their role in schwannomatosis-related pain (Figure 2). Of these genes the bradykinin receptor BDKRB1, was upregulated in DRGs following exposure to CM from painful tumors. This receptor is normally expressed at very low levels in sensory neurons but is induced and overexpressed in inflamed or damaged tissue by Il-1 ^27,28^. Interleukin 1β mRNA was also upregulated in DRG cells after treatment with CM from painful tumors. IL1 β is a proinflammatory cytokine that has been strongly implicated in nerve injury-induced tactile allodynia and in the development of chronic pain ^29^. Another finding of interest was the upregulation of TRPA1 mRNA transcription induced by SWN CM from two painful tumors (CM 2 and CM 3). TRPA1 plays a role in mechanical allodynia following induction of inflammatory and neuropathic pain ^30–34^. TRPA1 is implicated in acute pain behavior as well as the transition from acute to chronic pain^33^, plays a role in the detection of noxious cold and cold hypersensitivity^35^, and is activated by inflammatory mediators ^30–32^. Cross talk between the chemokine receptors, bradykinin, and TRPA1, and TRPV1 has been described ^36^. Therefore we examined agonists of TRP channels, TRPV1 and TRPA1 to determine whether these pathways are involved in the effects of painful SWN CM.

We found that conditioned media from painful schwannomatosis tumors sensitized mouse DRG neurons to the TRPV1 agonist capsaicin. Sensitization to this agent was evident as either an increase in response size or an increase to the percentage of DRG cells that responded to capsaicin. A heterogeneous response was elicited by the different painful CMs. CM from tumors 1 and 2 caused an increase in the percentage of DRG giving a response to capsaicin compared to non-painful CM but the magnitude (area under the curve) was similar between samples. Tumor CM 3 caused an increase in the magnitude of response. Conditioned media from painful tumor 4 caused an increase a magnitude of response to capsaicin as well as an increase in the percentage of neurons responding (Figure 5). Interestingly TRPV1 was not upregulated by painful CM at the transcriptional level suggesting that TRPV1 sensitization occurred post-transcriptionally.

Sensitivity to cinnamaldehyde was increased by both of the CMs that also induced transcriptional upregulation of TRPA1 mRNA (Figures 2&4). CM 3 produced a significantly larger AUC than any of the painful or non-painful CMs, while CM 2 caused a larger percentage of DRG cells to respond to cinnamaldehyde (Figure 4). The 3 other CMs from painful tumors had no effect on DRG response to cinnamaldehyde.

In our pursuit to identify substances from painful tumors which elicit effects on DRG neurons, we quantified cytokines secreted into CM. It is well known that Schwann cells secrete cytokines in models of nerve damage for myelin clearance and macrophage recruitment. Cytokine release may also contribute to a painful phenotype after injury ^10^. We found similar amounts of Il-6, CCL2/MCP-1, MIP-3a/CCL20, SERPINE1, PLAUR, CXCL5, DKK-1, and OPN in conditioned media from both painful and non-painful tumors (Supplemental Figure 3). These cytokines are known contributors to nociceptor sensitization after injury ^10,36–38^, but cannot account for the differential functional effects of CM from painful vs. nonpainful SWN tumors. Rather, we found increases in the levels of several cytokines secreted into the CM of painful tumors. One common cytokine, Growth/Differentiation Factor 15 (GDF-15/MIC-1), was found at increased levels in CM from all painful tumors but not in non-painful tumors (Figure 3). GDF-15 is a pro-survival factor for DRG neurons ^39^. GDF-15 has not been directly associated with pain but is secreted by Schwann cells after nerve injury ^40^. It is involved in peripheral nerve regeneration as it is shown to be axonally transported in lesioned peripheral nerves ^40^ and accelerates sensory recovery after injury ^41^. Levels of specific cytokines varied between the painful tumors. We found increases in the amounts of CD147, GM-CSF, CCL5/RANTES, VEGF, and IL-8 in CM from several painful tumors compared to the non-painful CM (Figure 3 and Table 1). Such a result is not unexpected, since the pain phenotypes observed among human patients with SWN are heterogeneous in nature. CD147, which was secreted by painful tumors is highly expressed in Schwann Cells ^42^. and is involved in the early stages of Schwann cell-axon contact. Secreted CD147 is an inducer of extracellular matrix metalloproteinases in peripheral nerve sheath tumors leading to progression of neurofibroma to Malignant Peripheral Nerve Sheath Tumors ^43,44^ It is upregulated in the extracellular matrix of DRGs after injury and is involved in mechanical allodynia in injured rats ^45^. Knockout of CD147 in mice causes attenuation of response to irritating odors^46^ and increases pain response to foot shock tests^47^.

**Table 1:** Levels of several cytokines are increased in CM from painful tumors Changes in the levels of 12 cytokines were detected in CM isolated painful and non-painful tumor cells. Increased levels of **CD147, GDF-15, GM-CSF, CXCL1**, and **PTX-3** were found in CM from painful tumor cell lines. Increased levels of IL-8 were detected in CM from painful tumors 3 and 4, **CCL5/RANTES** were noted in CM from painful tumor 2, increased levels of VEGF were found in CM from painful tumors 1 and 2, while increased levels of **angiogenin** were found in tumors from Painful tumors 1 and 3.

| CYTOKINE | CM 1 | CM 2 | CM 3 | CM 4 |
| --- | --- | --- | --- | --- |
| ANGIOGENIN | UP |  | UP |  |
| CD147 | UP | UP |  | UP |
| ENDOGLIN |  | UP |  |  |
| GDF-15/MIC-1 | UP | UP | UP | UP |
| GM-CSF | UP | UP |  | UP |
| CXCL1 | UP | UP | UP | UP |
| PTX3 | UP | UP | UP | UP |
| VEGF | UP | UP |  |  |
| IL8 |  |  | UP | UP |
| IL6 | UP | UP |  |  |
| RANTES/CCL5 |  | UP |  |  |
| G-CSF |  | UP |  | UP |

While GDF-15, and CD147 have implied functions in relation to pain, GM-CSF, CCL5/RANTES, IL-8 and VEGF, have been previously studied in signaling pathways that are related to pain response. GM-CSF contributes to some forms of tumor related pain by activating signaling cascades in DRG neurons, resulting in neuronal sensitization and increased membrane levels of TRPV1 ^48^. GM-CSF receptors are expressed in a subset of small/medium diameter neurons that also express TRPV1. Capsaicin responses can be increased after a 10 min incubation with GM-CSF, suggesting an interplay between receptors ^26^.

There is increasing evidence of crosstalk between receptors contributing to increases in intracellular calcium. One such example is VEGFR and TRPV1. VEGF sensitizes peripheral nociceptive neurons through actions on VEGFR2 and a TRPV1-dependent mechanism, enhancing nociceptive signaling ^49^. Both tumor CMs that caused a significant increase in the magnitude of response to capsaicin contained elevated levels of VEGF.

Co-activation of TRPA1 and TRPV1 has also been described ^50^. CM from painful tumor 2 sensitizes DRG neurons to capsaicin and cinnamaldehyde (Figures 4&5). A significant increase of CCL5/RANTES was found in the CM from Painful Tumor 2, but not that from other tumors (Figure 3, Supplemental Figure 4). It was recently demonstrated that CCL5 administration to the footpad of a mouse caused thermal hyperalgesia that could be prevented by co-administration of TRPV1 or TRPA1 inhibitors ^22^ In addition, upregulation of Bradykinin receptor, and TRPA1 in DRGs by conditioned media from painful tumors, suggests a potential interplay of signaling pathways leading to DRG hyper-excitability caused by factors released from tumors. BK receptor induces TRPA1 trafficking to the plasma membrane via PLC and PKA^51^. Cross talk between TRPV1, TRPA1, and Bradykinin receptor has been suggested in the development of chronic inflammatory pain ^36^.

Cultured DRG neurons express numerous types of chemokine receptors ^52^. One chemokine receptor that caught our attention is CXCR1/2. The ligand for this receptor is IL-8, which we found to be elevated in CM from painful tumors. IL-8 was secreted at high levels into the CMs that induced increased capsaicin sensitivity (CM 3 and 4, Figure 5, Table 1, Supplemental Figure 5). The receptors for IL-8 are expressed on DRG neurons and regulate the excitability of primary nociceptive neurons ^53^ and are implicated in chemotherapy induced neuropathic pain ^54^, as well as chronic inflammatory pain^55^. IL-8 is involved in long lasting mechanical hypersensitivity that persists even after the inflammatory response has resolved ^55^. Interestingly, IL-8 binding to receptors CXCR1/CXCR2 initiates a signaling cascade that induces transcription of NF-κb ^56^. In addition to IL-8, another SWN secreted protein, GM-CSF, activates NF-κb signaling through the GM-CSF receptor ^57^. NF-κb plays a role in persistent inflammatory pain ^58^. In a model of chronic inflammatory pain, NF-κB is translocated to the nucleus where it regulates a large number of genes and processes^58^. NF-κB is activated by several pain related inducers and at the same time regulates the expression of additional inducers, for example, Il-1β and the bradykinin receptor (which are transcriptionally upregulated by painful SWN CM in DRG neurons). NF-κB induces bradykinin receptor transcription in an inflammatory pain model ^59^. This suggests that multiple factors secreted by painful SWN CMs may influence or are influenced by NF-κB signaling in treated DRG neurons.

Collectively, the data presented in our study identify compelling candidate mechanisms by which some SWN tumors, but not others, produce pain. We have demonstrated that CM from painful schwannomas exhibit a differential capacity to sensitize DRG neurons, relative to CM from nonpainful tumors, and that this functional effect is associated with alterations in the expression of several pain-related target genes in the DRG. We also identified several secreted cytokines that may contribute to this effect. Finally, we have obtained evidence that the mechanisms that drive pain in SWN might be heterogeneous between tumors. Multiple cytokines and chemokines are secreted by painful schwannomas. These cytokines and chemokines can activate receptors in DRG neurons. Many of the receptors involved have the ability to activate other pain-related receptors by recruitment to the cell membrane or through cross-talk. Therefore, many players may be involved in the enhancement of neuronal sensitization in SWN patients. Further experimentation will be necessary to verify the specific involvement of these mechanisms in SWN-associated pain and identify additional pathways and players involved in this deleterious clinical process.

## Methods

### Cells and media

Schwannomatosis cell lines and conditioned media. Schwannomatosis-related tumors were collected from surgical cases from January 2014 through March 2017, occurring at Johns Hopkins School of Medicine. The diagnosis of Schwannomatosis was confirmed by Pathology. Informed, written consent was obtained prior to surgery. The Institutional Review Board (IRB) of the Johns Hopkins School of Medicine approved consent forms and study design. All research was performed in accordance with relevant guidelines/regulations and approved by the IRB. Schwannomatosis cell lines were established as described ^17^. Normal human Schwann cells were obtained from Sciencell (Carlsbad, Ca) Schwannomatosis or Schwann cells were cultured in DMEM containing 10% FBS and 2μM forskolin. When 60% confluence was reached, the medium was replaced with fresh medium. Conditioned medium (CM) was collected after 48 hours of incubation, centrifuged to remove cell debris and passed through a 0.22 micron filter. CM was frozen in aliquots for future use.

### Mouse Dorsal Root Ganglion (DRG) cultures

All experimental procedures were approved by the Institutional Animal Care and Use Committee of Johns Hopkins University School of Medicine and were in accordance with the guidelines provided by the National Institute of Health and the International Association for the Study of Pain. Isolation of dorsal root ganglion (DRG) neurons and primary culture were conducted as described ^60^. Briefly, DRG were isolated from 6-12 week old C57Bl6 wild type mice, rinsed in Complete Saline Solution (CSS containing, in mM: 137 NaCl, 5.3 KCl, 1 MgCl_2_, 3 CaCl_2_, 25 sorbitol, and 10 mM HEPES pH 7.2) and digested with TM Liberase (0.35 U/ml Roche) in in CSS/0.5mM EDTA at 37°C for 20 min, followed by TL Liberase (0.25 U/ml, Roche) in CSS/0.5mM EDTA supplemented with papain (30 U/ml, Worthington) for 15 min, centrifuged and resuspended in DMEM containing BSA (1 mg/ml, Sigma) and Trypsin Inhibitor (1 mg/ml, Sigma), and mechanically dissociated. Cells were passed through a 70 micron strainer and spotted on poly-l-lysine/laminin coated coverslips. After 1 hr. of adherence, complete DRG medium (DMEM/F12, 10% FBS, 1% glutamine, antibiotics), was added to each well.

### CM treatment of DRG neurons in culture

The day after establishment of primary mouse DRG cultures, culture medium was replaced with Schwannomatosis cell CM and the cells were further incubated for 48 hours at 37°C.

### Calcium imaging

Three separate DRG cultures were prepared for each Schwann cell CM to be tested. Four coverslips containing at least 30 neurons/slip were treated with CM from each Schwann cell line. A minimum of 100 DRG neurons were tested per condition. Cells were treated with CM for 48 hours before loading with 2 μM fura-2 acetoxymethyl ester (Molecular Probes) in calcium imaging buffer (CIB, containing in mM: 130 NaCl, 3 KCl, 2.5 CaCl2, 0,6 MgCl2, 10 HEPES, 1.2 NaHCO3, 10 glucose, pH 7.45, 290 mOsm adjusted with mannitol). Coverslips with fura-2-loaded cells were mounted on an inverted fluorescence microscope (TE200, Nikon). Images were acquired with a cMOS camera (NEO, Andor) using an excitation filter wheel (Ludl) equipped with 340 and 380 nm filters. Data were acquired using NIS Elements imaging software (Nikon). Fluorescence changes are expressed as the ratio of fluorescence emission at 520 nm upon stimulation at 340 to that upon stimulation at 380 nm (F340/F380). KCl experiments: A dose response curve of KCl was used to characterize overall neuronal sensitivity. KCl depolarizes the cell membrane, causing neurons to fire action potentials and triggering the opening of voltage-gated calcium channels, with a consequent rise in intracellular calcium. During continual imaging at ~2 s intervals, cell were successively exposed to modified CIB solutions supplemented with additional KCl (at net concentrations of 10 mM, 15 mM, 20 mM, or 25 mM, with equimolar removal of NaCl) delivered by perfusion for 30 s each, with a recovery period of washing with unmodified CIB (containing 3 mM KCl) for 2 min between concentrations. Capsaicin experiments: Capsaicin was dissolved in CIB at a net concentration of 25nM. During continual imaging at ~2 sec intervals cells were perfused with 25nM Capsaicin for 30 seconds with a recovery period of washing with unmodified CIB. Cinnamaldehyde experiments: Similarly, DRG cells were exposed with 200 uM cinnamaldehyde/CIB solution for 4 minutes followed by a 4 minute washout with CIB.

### RNA extraction, cDNA, RT^2^ Profiler PCR Arrays

DRG neurons were harvested after 48 hours exposure to CM and total RNA was extracted using Trizol (Invitrogen). 250 ng of total RNA was converted to cDNA using the RT2 Reverse Strand kit (Qiagen). The mouse RT^2^ neuropathic and inflammatory PCR array (Qiagen) was used for qPCR according to the manufacturer’s instructions. In brief, 250 ng of cDNA was mixed with RT^2^ SYBR Green qPCR Mastermix and water. 25 μl of DNA/enzyme/buffer mix was pipetted into each well of the supplied 96 well plate. Amplifications were carried out in 96 well plates in a Roche Lightcycler 480 with cycling as follows: 95 °C 10 min (hot start), and 45 cycles of 95 °C 15 sec and 60°C for 1 min. All reactions were performed in triplicate. Dissociation curve analysis was performed to rule out experimental PCR artifacts or non-specific amplification. Expression of genes relative to glyceraldehyde-3-phosphate dehydrogenase (GAPDH) and β2 microglobulin (β 2M) was calculated based on the threshold cycle (Ct) as 2-Δ(ΔCt), where ΔCt = Ct, GENE – average Ct for GAPDH and β2M, and Δ(ΔCt) = ΔCt P-ΔCt NP (P, painful tumor, NP, nonpainful tumor.) p values were calculated by Student’s t-test https://dataanalysis.qiagen.com/pcr/arrayanalysis

### Cytokine arrays

CM was examined for the presence of secreted cytokines using the Proteome Profiler Human Cytokine XL Array (R&D Systems) according to the manufacturer’s instructions. This array detects 102 cytokines simultaneously on a single membrane using pre-spotted capture antibodies. 500 microliters of CM were run on each membrane in separate dishes (Non-painful Tumors, Painful Tumor 1, Painful Tumor 2, Painful Tumor 3, Painful Tumor 4).The captured target proteins from conditioned media were detected with biotinylated detection antibodies and then visualized using chemiluminescent detection reagents with a 2 minute exposure to X-ray film.

### Data analysis and Statistics

For cytokine arrays, X-ray films were scanned using a gel imager and quantified using ImageJ. Pixel densities were compared and averaged for the duplicate spots on the blots, normalized to the standard provided, and plotted using Prism 8. For qRT-PCR, Fold change in gene expression was determined by the 2^(- Delta Delta Ct) method following the guidelines for the RT2 Profiler PCR Array Data Analysis Webportal for cataloged RT2 Profiler array (Neuropathic and Inflammatory PCR array PAMM-162Z). Statistical significance of the RT2 PCR data was determined using p values were calculated by Student’s t-test (https://dataanalysis.qiagen.com/pcr/arrayanalysis). Standard error (SE) of mean was also calculated for the control and test groups as (SE= SD/√n-1), where SD is standard deviation and n is number of samples. For calcium imaging, area under the curve of fura-2 ratio, with subtraction of the baseline obtained prior to each stimulus, was used to compare the effects of painful vs. non-painful CM. For ease of presentation, data for the non-painful tumors (n=3) were averaged together. The differences in AUC between non-painful CMs were not significant (Supplemental figure 2). ANOVA analysis with Dunnet multiple comparisons was used to compare the effects of CM groups (Graphpad Prism8). In addition, the percentage of cells responding to KCl, capsaicin, or cinnamaldehyde stimulation was analyzed. An individual cell’s response to stimuli was considered to be positive if the fura-2 ratio was greater than 0.1 after baseline subtraction. Chi-squared analysis was performed to test significance between the percentage of cells responding to painful and non-painful CM. The authors will make materials, data and protocols available upon request.

## Acknowledgements

Dr. Miriam and Sheldon Adelson Medical Foundation, the Ohrstrom Foundation, the Pamela Mars Wright Foundation, and The Johns Hopkins Neurosurgery Pain Research Institute supported this work. We thank members of the Hoke and Caterina labs for helpful discussions and Randy Rubright for technical support.

## Author Contributions

K.L.O. conceived the project, designed the experiments, conducted experiments, analyzed the data, and wrote the manuscript. K.J.D conducted experiments, performed data analysis, and reviewed the manuscript. A.B. M.J.C. and A.H. facilitated experimental design, assisted in data analysis, and revised the manuscript.

## Additional information

Disclosure: MJC was previously on the Scientific Advisory Board of Hydra Biosciences, which works on TRPA1 related products. The Johns Hopkins Office of Policy Coordination is managing this potential/perceived conflict.

## References

1 MacCollin, M. et al. Diagnostic criteria for schwannomatosis. Neurology 64, 1838–1845, doi:10.1212/01.WNL.0000163982.78900.AD (2005).

2 Merker, V. L. et al. Relationship between whole-body tumor burden, clinical phenotype, and quality of life in patients with neurofibromatosis. Am J Med Genet A 164A, 1431–1437, doi:10.1002/ajmg.a.36466 (2014).

3 Merker, V. L., Esparza, S., Smith, M. J., Stemmer-Rachamimov, A. & Plotkin, S. R. Clinical features of schwannomatosis: a retrospective analysis of 87 patients. Oncologist 17, 1317–1322, doi:10.1634/theoncologist.2012-0162 (2012).

4 Blakeley, J. et al. Clinical response to bevacizumab in schwannomatosis. Neurology 83, 1986–1987, doi:10.1212/WNL.0000000000000997 (2014).

5 Gonzalvo, A. et al. Schwannomatosis, sporadic schwannomatosis, and familial schwannomatosis: a surgical series with long-term follow-up. Clinical article. J Neurosurg 114, 756–762, doi:10.3171/2010.8.JNS091900 (2011).

6 Huang, J. H., Simon, S. L., Nagpal, S., Nelson, P. T. & Zager, E. L. Management of patients with schwannomatosis: report of six cases and review of the literature. Surg Neurol 62, 353–361; discussion 361, doi:10.1016/j.surneu.2003.11.020 (2004).

7 Halvorsen, C. M. et al. The Long-term Outcome After Resection of Intraspinal Nerve Sheath Tumors: Report of 131 Consecutive Cases. Neurosurgery 77, 585–592; discussion 592–583, doi:10.1227/NEU.0000000000000890 (2015).

8 Mansukhani, S. A., Butala, R. P., Shetty, S. H. & Khedekar, R. G. Familial Schwannomatosis: A Diagnostic Challenge. J Clin Diagn Res 11, RD01–RD03, doi:10.7860/JCDR/2017/20929.9307 (2017).

9 MacCollin, M., Woodfin, W., Kronn, D. & Short, M. P. Schwannomatosis: a clinical and pathologic study. Neurology 46, 1072–1079 (1996).

10 Campana, W. M. Schwann cells: activated peripheral glia and their role in neuropathic pain. Brain BehavImmun 21, 522–527, doi:10.1016/j.bbi.2006.12.008 (2007).

11 Okamoto, K., Martin, D. P., Schmelzer, J. D., Mitsui, Y. & Low, P. A. Pro-and anti-inflammatory cytokine gene expression in rat sciatic nerve chronic constriction injury model of neuropathic pain. Exp Neurol 169, 386–391, doi:10.1006/exnr.2001.7677 (2001).

12 Ozaki, A., Nagai, A., Lee, Y. B., Myong, N. H. & Kim, S. U. Expression of cytokines and cytokine receptors in human Schwann cells. Neuroreport 19, 31–35, doi:10.1097/WNR.0b013e3282f27e60 (2008).

13 Be’eri, H., Reichert, F., Saada, A. & Rotshenker, S. The cytokine network of wallerian degeneration: IL-10 and GM-CSF. Eur J Neurosci 10, 2707–2713 (1998).

14 Shamash, S., Reichert, F. & Rotshenker, S. The cytokine network of Wallerian degeneration: tumor necrosis factor-alpha, interleukin-1alpha, and interleukin-1beta. J Neurosci 22, 3052–3060, doi:20026249 (2002).

15 Watkins, L. R., Milligan, E. D. & Maier, S. F. Glial proinflammatory cytokines mediate exaggerated pain states: implications for clinical pain. Adv Exp Med Biol 521, 1–21 (2003).

16 Siqueira Mietto, B. et al. Role of IL-10 in Resolution of Inflammation and Functional Recovery after Peripheral Nerve Injury. J Neurosci 35, 16431–16442, doi:10.1523/JNEUROSCI.2119-15.2015 (2015).

17 Ostrow, K. L., Donaldson, K., Blakeley, J., Belzberg, A. & Hoke, A. Immortalized Human Schwann Cell Lines Derived From Tumors of Schwannomatosis Patients. PLoS One 10, e0144620, doi:10.1371/journal.pone.0144620 (2015).

18 D’Arco, M., Margas, W., Cassidy, J. S. & Dolphin, A. C. The upregulation of alpha2delta-1 subunit modulates activity-dependent Ca2+ signals in sensory neurons. JNeurosci 35, 5891–5903, doi:10.1523/JNEUROSCI.3997-14.2015 (2015).

19 Hall, K. E., Sima, A. A. & Wiley, J. W. Opiate-mediated inhibition of calcium signaling is decreased in dorsal root ganglion neurons from the diabetic BB/W rat. J Clin Invest 97, 1165–1172, doi:10.1172/JCI118530 (1996).

20 Kim, Y. S. et al. Central terminal sensitization of TRPV1 by descending serotonergic facilitation modulates chronic pain. Neuron 81, 873–887, doi:10.1016/j.neuron.2013.12.011 (2014).

21 Teichert, R. W. et al. Functional profiling of neurons through cellular neuropharmacology. Proc Natl Acad Sci U S A 109, 1388–1395, doi:10.1073/pnas.1118833109 (2012).

22 Gonzalez-Rodriguez, S. et al. Hyperalgesic and hypoalgesic mechanisms evoked by the acute administration of CCL5 in mice. Brain Behav Immun 62, 151–161, doi:10.1016/j.bbi.2017.01.014 (2017).

23 Kimball, E. S. et al. Stimulation of neuronal receptors, neuropeptides and cytokines during experimental oil of mustard colitis. Neurogastroenterol Motil 19, 390–400, doi:10.1111/j.1365-2982.2007.00939.x (2007).

24 Caterina, M. J. et al. Impaired nociception and pain sensation in mice lacking the capsaicin receptor. Science 288, 306–313 (2000).

25 Caterina, M. J. et al. The capsaicin receptor: a heat-activated ion channel in the pain pathway. Nature 389, 816–824, doi:10.1038/39807 (1997).

26 Donatien, P. et al. Granulocyte-macrophage colony-stimulating factor receptor expression in clinical pain disorder tissues and role in neuronal sensitization. Pain Rep 3, e676, doi:10.1097/PR9.0000000000000676 (2018).

27 Couture, R., Harrisson, M., Vianna, R. M. & Cloutier, F. Kinin receptors in pain and inflammation. Eur J Pharmacol 429, 161–176 (2001).

28 Petcu, M. et al. Role of kinin B1 and B2 receptors in a rat model of neuropathic pain. Int Immunopharmacol 8, 188–196, doi:10.1016/j.intimp.2007.09.009 (2008).

29 Gui, W. S. et al. Interleukin-1beta overproduction is a common cause for neuropathic pain, memory deficit, and depression following peripheral nerve injury in rodents. Mol Pain 12, doi:10.1177/1744806916646784 (2016).

30 Bautista, D. M., Pellegrino, M. & Tsunozaki, M. TRPA1: A gatekeeper for inflammation. Annu Rev Physiol 75, 181–200, doi:10.1146/annurev-physiol-030212-183811 (2013).

31 Gouin, O. et al. TRPV1 and TRPA1 in cutaneous neurogenic and chronic inflammation: pro-inflammatory response induced by their activation and their sensitization. Protein Cell 8, 644–661, doi:10.1007/s13238-017-0395-5 (2017).

32 Green, D. et al. Central activation of TRPV1 and TRPA1 by novel endogenous agonists contributes to mechanical allodynia and thermal hyperalgesia after burn injury. Mol Pain 12, doi:10.1177/1744806916661725 (2016).

33 Kadkova, A., Synytsya, V., Krusek, J., Zimova, L. & Vlachova, V. Molecular basis of TRPA1 regulation in nociceptive neurons. A review. Physiol Res 66, 425–439 (2017).

34 Koivisto, A. et al. TRPA1: a transducer and amplifier of pain and inflammation. Basic Clin Pharmacol Toxicol 114, 50–55, doi:10.1111/bcpt.12138 (2014).

35 del Camino, D. et al. TRPA1 contributes to cold hypersensitivity. J Neurosci 30, 15165–15174, doi:10.1523/JNEUROSCI.2580-10.2010 (2010).

36 White, F. A., Jung, H. & Miller, R. J. Chemokines and the pathophysiology of neuropathic pain. Proc Natl Acad Sci U S A 104, 20151–20158, doi:10.1073/pnas.0709250104 (2007).

37 Wei, X. H. et al. The up-regulation of IL-6 in DRG and spinal dorsal horn contributes to neuropathic pain following L5 ventral root transection. Exp Neurol 241, 159–168, doi:10.1016/j.expneurol.2012.12.007 (2013).

38 Zhang, Z. J., Jiang, B. C. & Gao, Y. J. Chemokines in neuron-glial cell interaction and pathogenesis of neuropathic pain. Cell Mol Life Sci 74, 3275–3291, doi:10.1007/s00018-017-2513-1 (2017).

39 Strelau, J., Schober, A., Sullivan, A., Schilling, L. & Unsicker, K. Growth/differentiation factor-15 (GDF-15), a novel member of the TGF-beta superfamily, promotes survival of lesioned mesencephalic dopaminergic neurons in vitro and in vivo and is induced in neurons following cortical lesioning. J Neural Transm Suppl, 197–203 (2003).

40 Strelau, J. et al. Progressive postnatal motoneuron loss in mice lacking GDF-15. J Neurosci 29, 13640–13648, doi:10.1523/JNEUROSCI.1133-09.2009 (2009).

41 Mensching, L. et al. Local substitution of GDF-15 improves axonal and sensory recovery after peripheral nerve injury. Cell Tissue Res 350, 225–238, doi:10.1007/s00441-012-1493-6 (2012).

42 Spiegel, I. et al. Identification of novel cell-adhesion molecules in peripheral nerves using a signal-sequence trap. Neuron Glia Biol 2, 27–38, doi:10.1017/S1740925X0600007X (2006).

43 Zhu, X., Song, Z., Zhang, S., Nanda, A. & Li, G. CD147: a novel modulator of inflammatory and immune disorders. Curr Med Chem 21, 2138–2145 (2014).

44 Nabeshima, K. et al. Expression of emmprin and matrix metalloproteinases (MMPs) in peripheral nerve sheath tumors: emmprin and membrane-type (MT)1-MMP expressions are associated with malignant potential. Anticancer Res 26, 1359–1367 (2006).

45 Wang, Q. et al. Upregulation of EMMPRIN (OX47) in Rat Dorsal Root Ganglion Contributes to the Development of Mechanical Allodynia after Nerve Injury. Neural Plast 2015, 249756, doi:10.1155/2015/249756 (2015).

46 Igakura, T. et al. Roles of basigin, a member of the immunoglobulin superfamily, in behavior as to an irritating odor, lymphocyte response, and blood-brain barrier. Biochem Biophys Res Commun 224, 33–36, doi:10.1006/bbrc.1996.0980 (1996).

47 Naruhashi, K. et al. Abnormalities of sensory and memory functions in mice lacking Bsg gene. Biochem Biophys Res Commun 236, 733–737, doi:10.1006/bbrc.1997.6993 (1997).

48 Schweizerhof, M. et al. Hematopoietic colony-stimulating factors mediate tumor-nerve interactions and bone cancer pain. Nat Med 15, 802–807, doi:10.1038/nm.1976 (2009).

49 Turker, E. et al. Vascular Endothelial Growth Factor (VEGF) Induced Downstream Responses to Transient Receptor Potential Vanilloid 1 (TRPV1) and 3-Iodothyronamine (3-T1AM) in Human Corneal Keratocytes. Front Endocrinol (Lausanne) 9, 670, doi:10.3389/fendo.2018.00670 (2018).

50 Ruparel, N. B., Patwardhan, A. M., Akopian, A. N. & Hargreaves, K. M. Homologous and heterologous desensitization of capsaicin and mustard oil responses utilize different cellular pathways in nociceptors. Pain 135, 271–279, doi:10.1016/j.pain.2007.06.005 (2008).

51 Lapointe, T. K. & Altier, C. The role of TRPA1 in visceral inflammation and pain. Channels (Austin) 5, 525–529, doi:10.4161/chan.5.6.18016 (2011).

52 Miller, R. J., Jung, H., Bhangoo, S. K. & White, F. A. Cytokine and chemokine regulation of sensory neuron function. Handb Exp Pharmacol, 417–449, doi:10.1007/978-3-540-79090-7_12 (2009).

53 Li, X. et al. Enhanced RAGE Expression in the Dorsal Root Ganglion May Contribute to Neuropathic Pain Induced by Spinal Nerve Ligation in Rats. Pain Med 17, 803–812, doi:10.1093/pm/pnv035 (2016).

54 Brandolini, L. et al. CXCR1/2 pathways in paclitaxel-induced neuropathic pain. Oncotarget 8, 23188–23201, doi:10.18632/oncotarget.15533 (2017).

55 Sachs, D., Cunha, F. Q., Poole, S. & Ferreira, S. H. Tumour necrosis factor-alpha, interleukin-1beta and interleukin-8 induce persistent mechanical nociceptor hypersensitivity. Pain 96, 89–97 (2002).

56 Waugh, D. J. & Wilson, C. The interleukin-8 pathway in cancer. Clin Cancer Res 14, 6735–6741, doi:10.1158/1078-0432.CCR-07-4843 (2008).

57 Ebner, K., Bandion, A., Binder, B. R., de Martin, R. & Schmid, J. A. GMCSF activates NF-kappaB via direct interaction of the GMCSF receptor with IkappaB kinase beta. Blood 102, 192–199, doi:10.1182/blood-2002-12-3753 (2003).

58 Souza, G. R. et al. Involvement of nuclear factor kappa B in the maintenance of persistent inflammatory hypernociception. Pharmacol Biochem Behav 134, 49–56, doi:10.1016/j.pbb.2015.04.005 (2015).

59 Passos, G. F. et al. Kinin B1 receptor up-regulation after lipopolysaccharide administration: role of proinflammatory cytokines and neutrophil influx. J Immunol 172, 1839–1847, doi:10.4049/jimmunol.172.3.1839 (2004).

60. Qu, L. & Caterina, M. J. Enhanced excitability and suppression of A-type K(+) currents in joint sensory neurons in a murine model of antigen-induced arthritis. Sci Rep 6, 28899, doi:10.1038/srep28899 (2016).

