## Supplemental Figures for "The Secretomes of Painful Versus Nonpainful Human Schwannomatosis Tumor Cells Differentially Influence Sensory Neuron Gene Expression and Sensitivity"

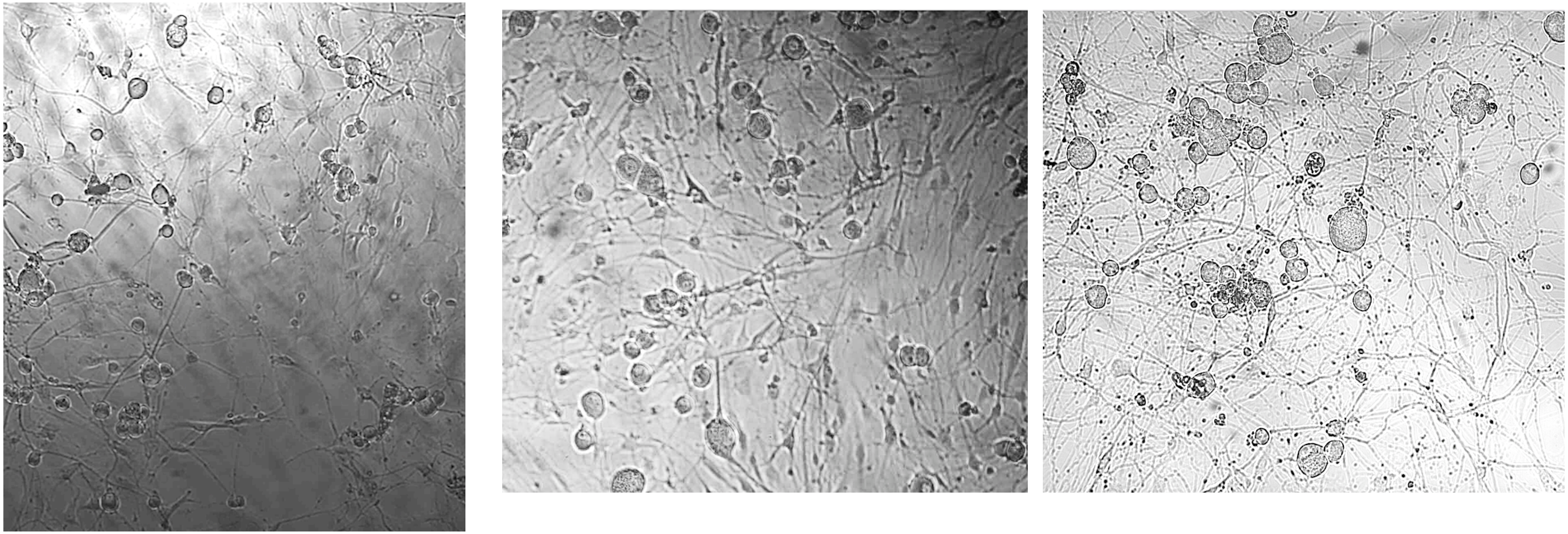

Painful Tumor 1 CM

Painful Tumor 2 CM

Non-Painful Tumor 2 CM

Supplemental Figure 1: Brightfield images of CM-treated DRGs prior to Calcium imaging. Neurite sprouting can be visualized in DRGs after 48 hours of incubation with SWN tumor CM regardless of pain status.

### Dose response to KCl After treatment with NP Tumor CM

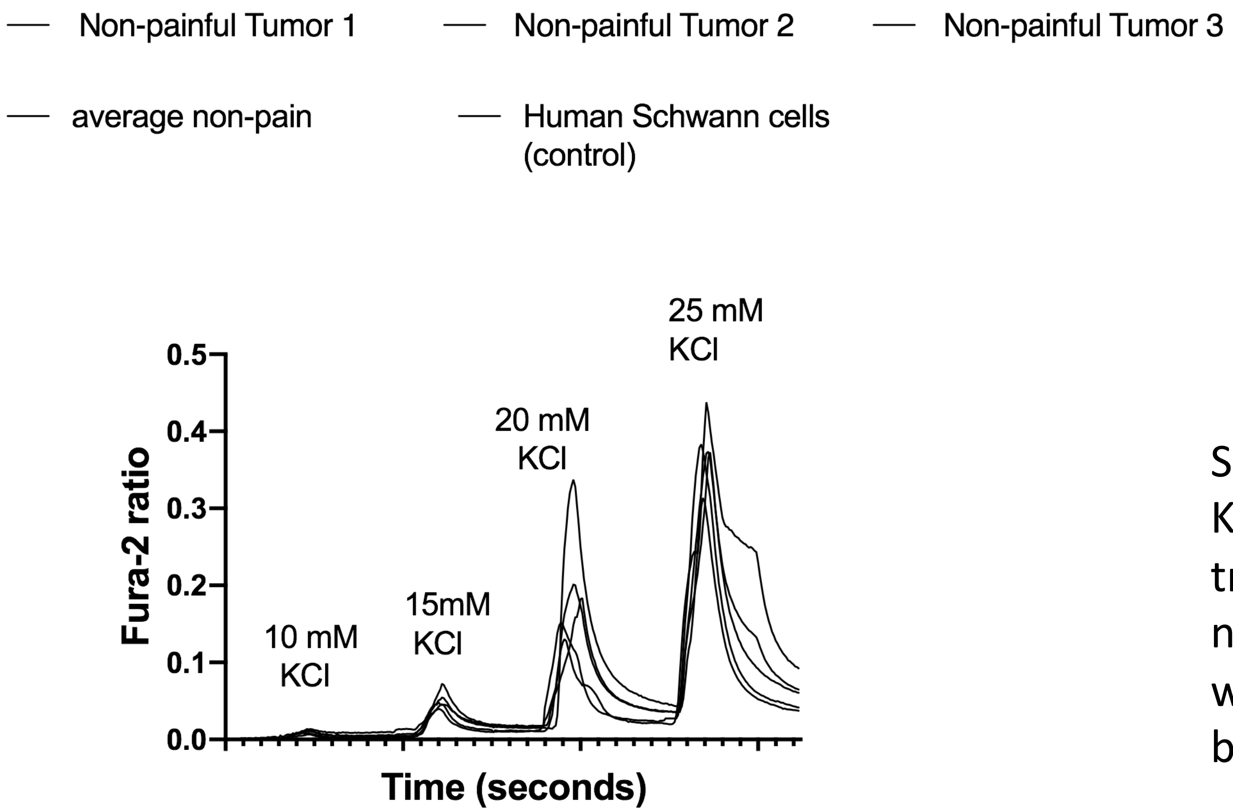


Supplemental Figure 2:
KCL dose response after pre-treatment with 3 separate non-painful tumor CMs. AUC was not statistically significant between groups.

| KCL Dose | Average AUC NP Tumor 1 | Average AUC NP Tumor 2 | Average AUC NP Tumor 3 | p value |
|---|---|---|---|---|
| 10 mM | 0.243 (+/- 0.07) | 0.120 (+/- 0.08) | 0.221 (+/- 0.06) | p=0.37 |
| 15 mM | 0.946 (+/- 0.12) | 0.85 (+/- 0.24) | 1.14 (+/- 0.16) | p=0.4 |
| 20 mM | 3.12 (+/- 0.51) | 2.86 (+/- 0.53) | 4.2 (+/- 0.59) | p=0.2 |
| 25 mM | 6.45 (+/- 1.68) | 6.92 (+/- 1.58) | 6.94 (+/- 0.74) | p-0.9 |

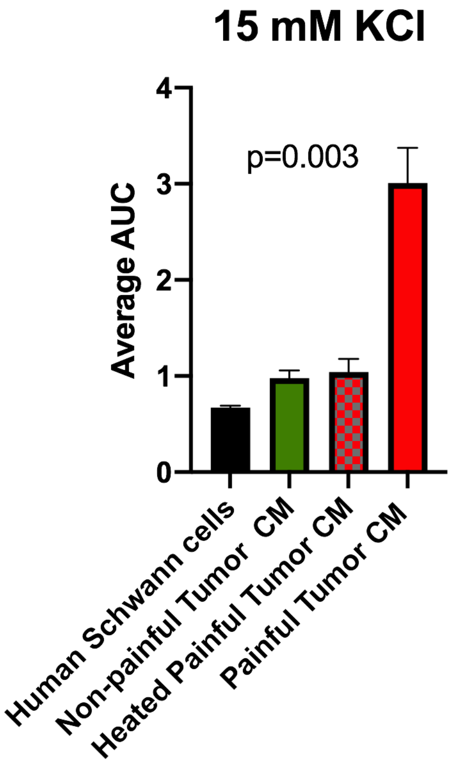


Supplemental Figure 3: Effect of Heating painful Tumor CM. Heating the painful CM at 80C for 30 min (**red patterned**) reduced responsiveness to that of non-painful CM (green)

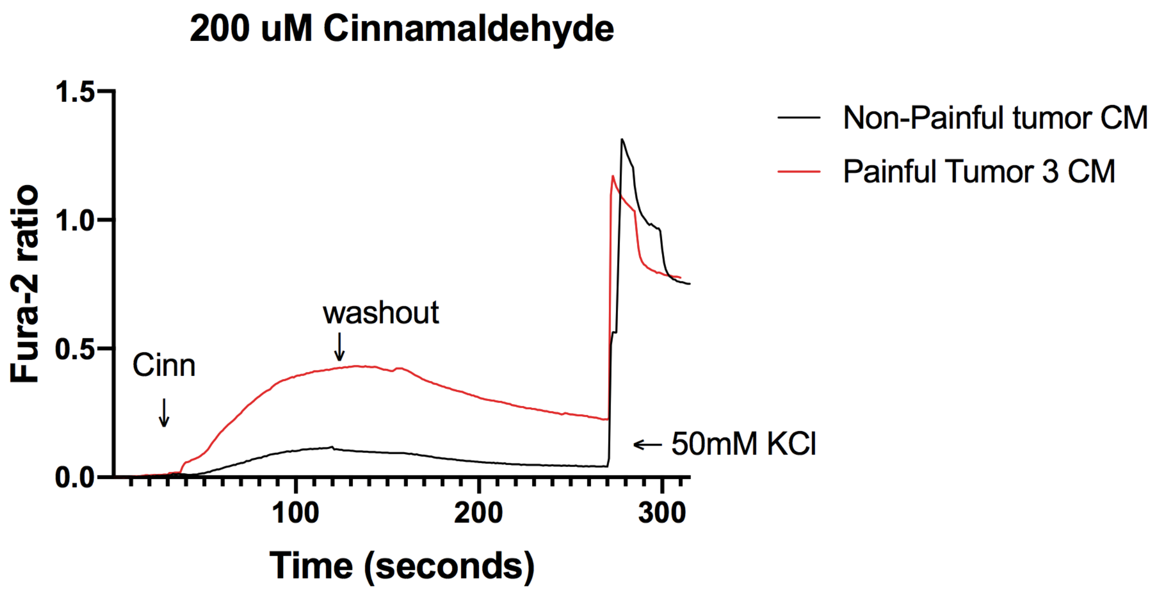


Supplemental Figure 4: DRG cells were perfused 200 uM Cinnamaldehyde for 120 sec with a washout with 3mM KCl buffer for and additional 2 minutes. Fura-2 ratio measurements were recorded at 2 second intervals. The datapoints were corrected for differing baseline readings of Fura-2. Area under the curve was calculated for each treatment. The effect of cinnamaldehyde started 30-60 seconds after treatment peaked at 120 seconds, and lasted for 4 minutes. Painful tumor 3 CM increased the responsiveness of the DRGs to cinnamaldehyde as calculated by area under the curve $p=0.03$.

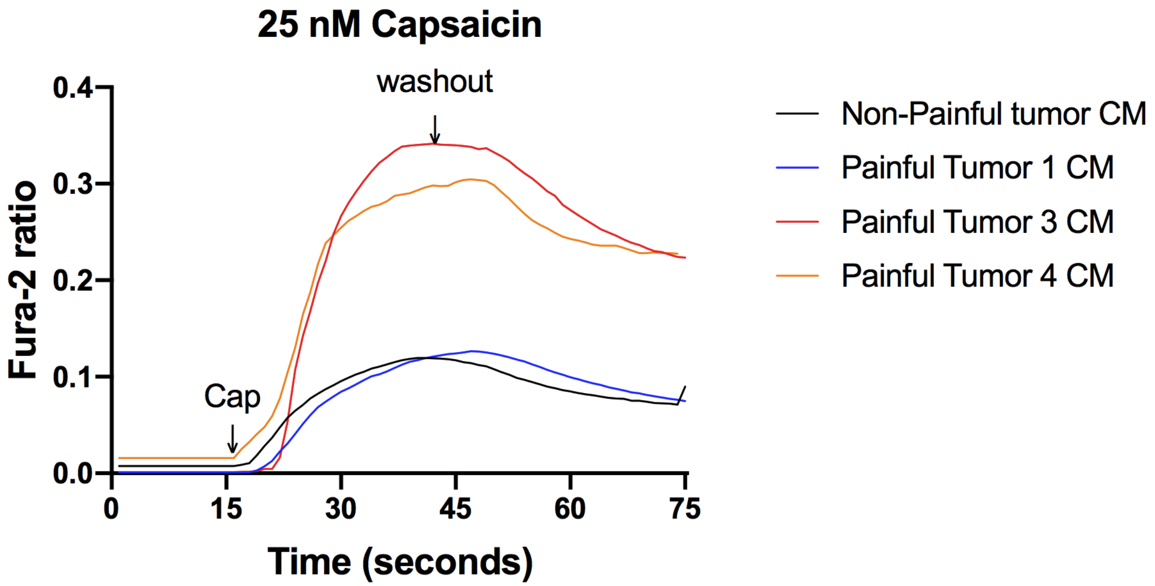


Supplemental Figure 5: DRG cells were perfused 25nM capsaicin for 30sec with a washout with 3mM KCl buffer for 2 minutes. Fura-2 ratio measurements were recorded at 2 second intervals. The datapoints were corrected for differing baseline readings of Fura-2. Area under the curve was calculated for each treatment. The effect of  capsaicin started immediately peaked after 30 seconds lasted for 1 minute. CM from painful tumors 3 and 4 CM increased the responsiveness of the DRGs to capsaicin as calculated by area under the curve p=0.0001. One way ANOVA with multiple comparisons was employed to determine statistical significance.

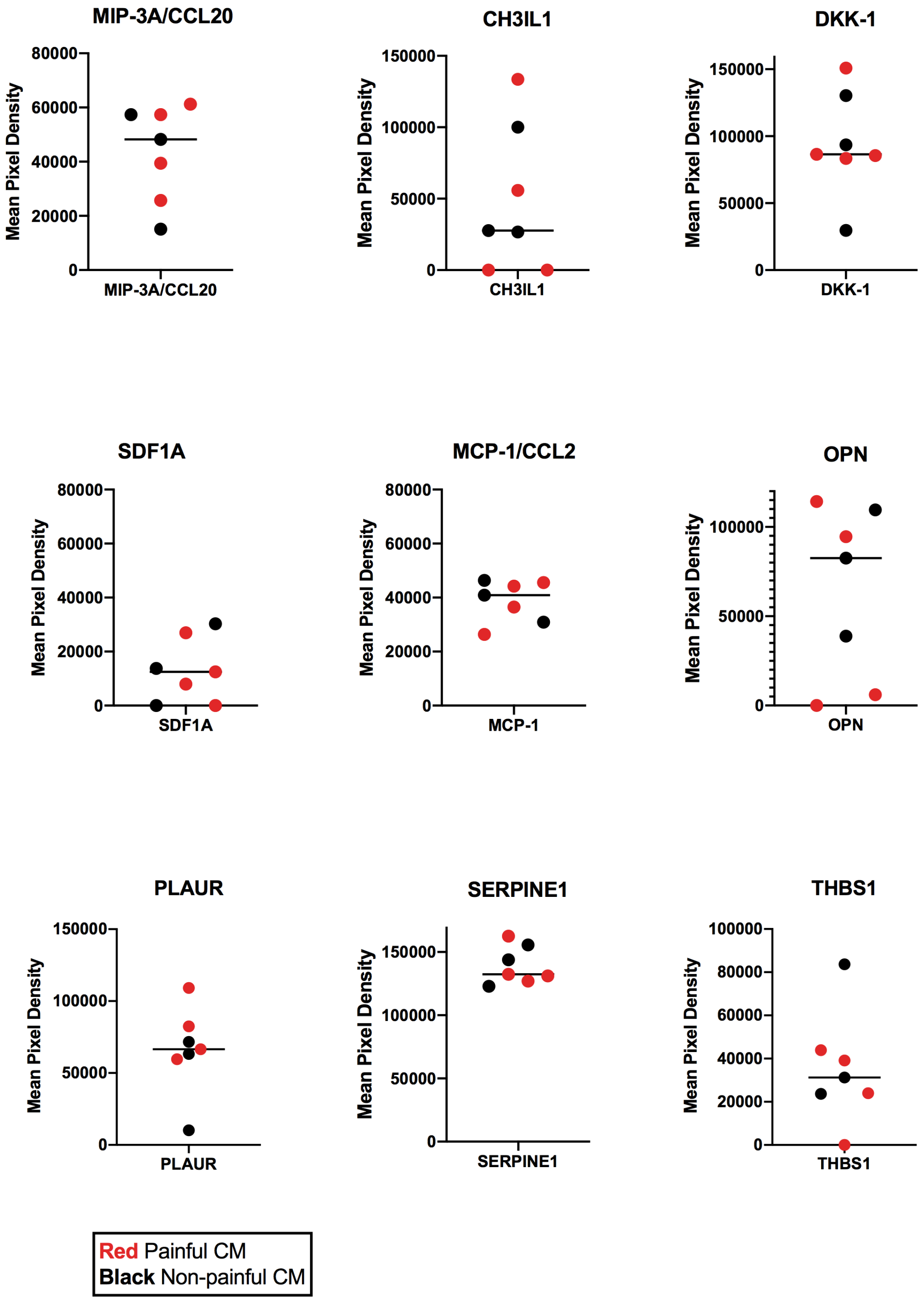


Supplemental Figure 6: Cytokines that were detected in CM that were similar between painful and Non-painful CM.

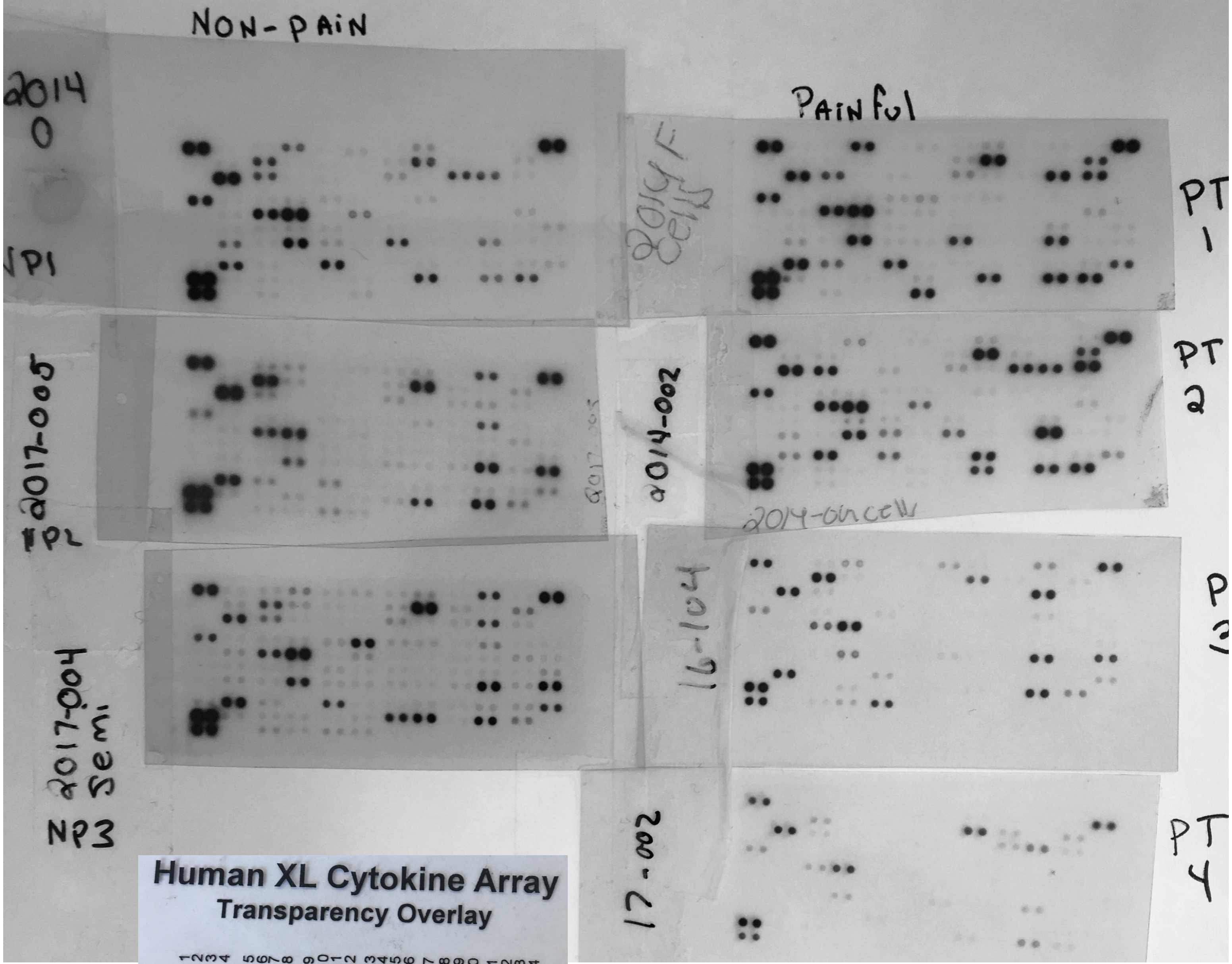


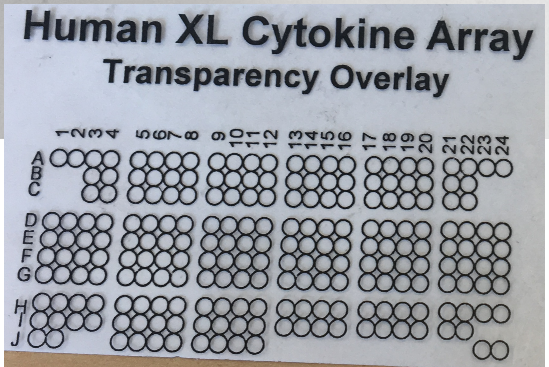


Supplemental Figure 7: Human cytokine array blots after 2 min exposure to X-ray film. Positions A 1,2 A23,24 and J 23, 24 are positive controls used for normalizing protein levels between blots.
